# Antibiotic-driven Escape of Host in a Parasite-induced Red Queen Dynamics

**DOI:** 10.1101/351940

**Authors:** Elizabeth L. Anzia, Jomar F. Rabajante

**Author notes:** **One sentence summary:** Uninterrupted intake of highly effective antibiotic can be a strategy to escape the Red Queen dynamics.

## Abstract

Winnerless coevolution of hosts and parasites could exhibit Red Queen dynamics, which is characterized by parasite-driven cyclic switching of expressed host phenotypes. We hypothesize that the application of antibiotics to suppress the reproduction of parasites can provide opportunity for the hosts to escape such winnerless coevolution. Here, we formulate a minimal mathematical model of host-parasite interaction involving multiple host phenotypes that are targeted by adapting parasites. Our model predicts the levels of antibiotic effectiveness that can steer the parasite-driven cyclic switching of host phenotypes (heteroclinic oscillations) to a stable equilibrium of host survival. Our simulations show that uninterrupted application of antibiotic with high-level effectiveness (> 85%) is needed to escape the Red Queen dynamics. Intermittent and low level of antibiotic effectiveness are indeed useless to stop host-parasite coevolution. This study can be a guide in designing good practices and protocols to minimize risk of further progression of parasitic infections.

## Introduction

Understanding antagonism-mediated evolution is essential as antagonistic interactions, including parasitism, play major roles in the formation and maintenance of the structure of communities [1, 2, 3, 4, 5]. The Red Queen hypothesis is a model for winnerless antagonistic coevolution between interacting species, such as host-parasite, prey-predator and victim-exploiter [6, 7, 8]. The Red Queen hypothesis has been demonstrated using various schemes, e.g., to explain the evolution of sex [9, 10, 11] and the antagonism-mediated species diversity [6, 12, 13]. Here, we focus on fluctuating Red Queen mode (in contrast to escalatory Red Queen and chase Red Queen [14, 9]) to explain the effect of inhibiting the growth of parasites on host-parasite coevolution. One of the numerical manifestations of fluctuating Red Queen mode is the canonical Red Queen dynamics/cycles [15, 16, 17].

The Red Queen dynamics in host-parasite system describes winnerless coevolution between hosts and parasites, realized through perpetual negative frequency-dependent selection [16]. The canonical case of the Red Queen dynamics follows the following process: host population (say, H1) evolves to a new type (H2) to escape their parasites (P1). However, the decline in the population of H1 will drive parasite population P1 to evolve to a new strain (P2) as a response against the evolution of the host. The new parasite P2 can infect H2, which will drive the hosts to evolve again, resulting in never-ending alternating cycles of dominance (Figure 1 and 2A). For example, the Red Queen dynamics explain coevolution in invertebrate-parasite systems, such as infection of *Daphnia magna* by bacteria *Pasteuria ramosa* [18, 19].

**Figure 1:**
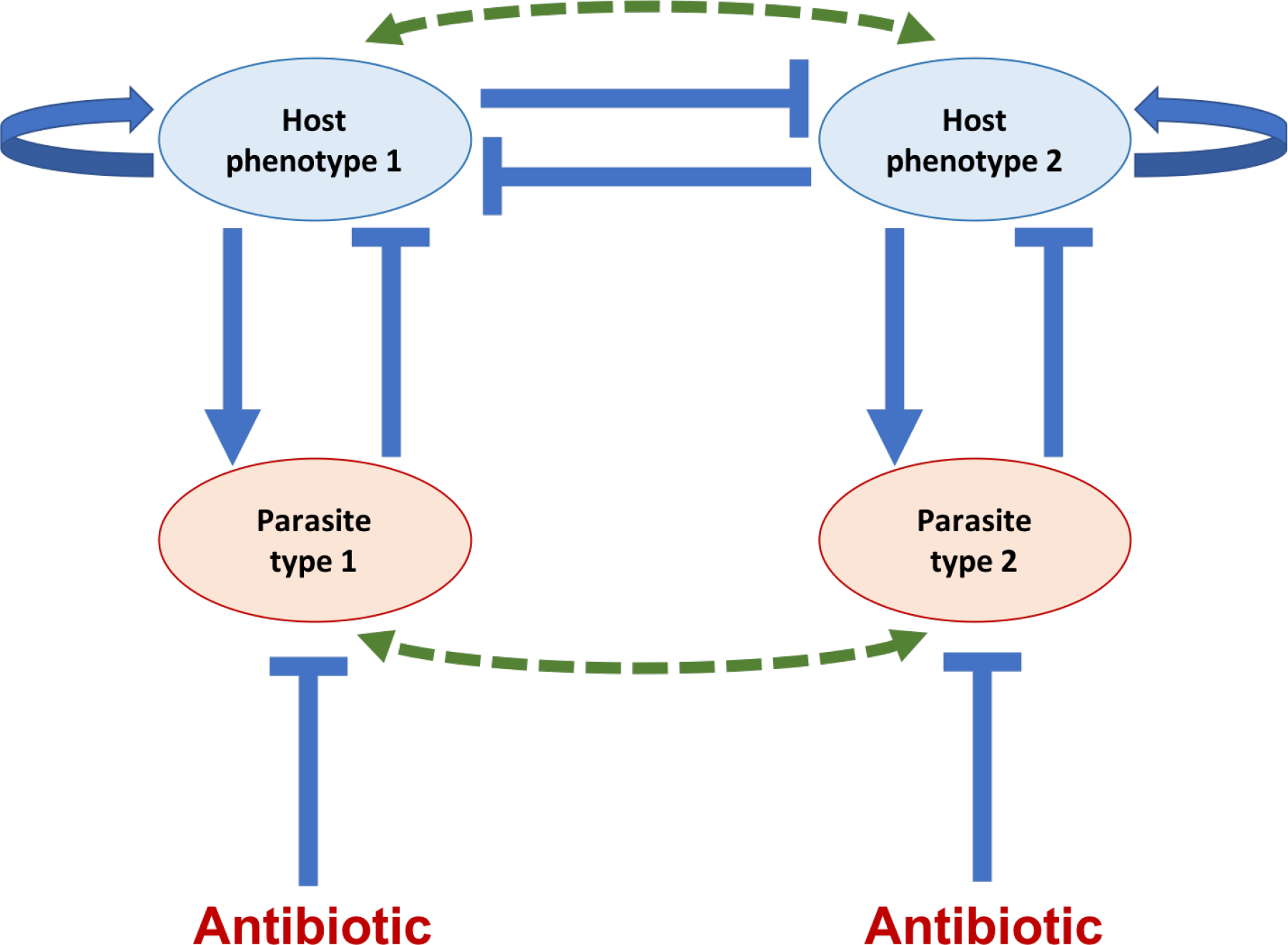
The host phenotype decision-switch network with the influence of para-sitism. Self-regulation in hosts means host growth and phenotypic memory. Host phenotypes mutually inhibit each other to characterize trait selection, but simula-tions can show coexistence is possible. Parasites proliferate through infection but their growth can be repressed by antibiotics. Solid arrows denote positive interac-tion. Solid bars denote negative interaction. Dashed arrows imply evolution.

**Figure 2:**
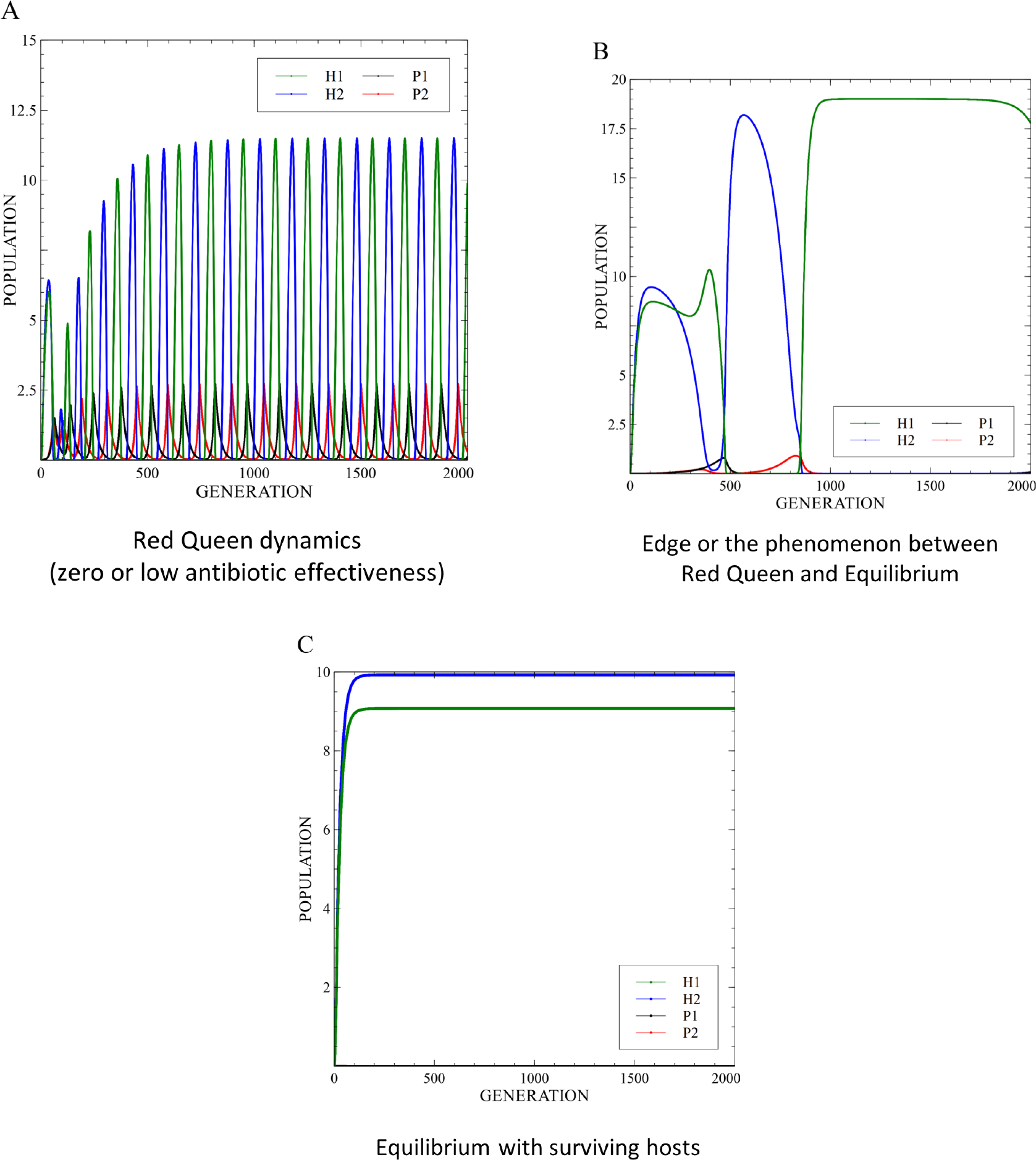
Different qualitative behaviors of host-parasite interaction with the addi-tional effect of antibiotics to parasite. (A) Oscillating population frequencies of hosts and parasites as they interact with each other. The effect of insufficient levels of an-tibiotics to eradicate parasitism can also result in Red Queen dynamics. (B) Irregular oscillating population frequencies of hosts and parasites due to the insufficient effect of antibiotics. (C) The population frequencies of hosts and parasites converge to an equilibrium because the levels of the antibiotics are sufficient to eradicate parasitism.

The canonical Red Queen dynamics can be characterized by populations of host types (and parasite types) undergoing cyclic dominance switching in population abundance, where every host (parasite) type has the opportunity to have abundant frequency for some period of time [16]. Mathematically, we define Red Queen dynamics as population oscillations that are out-of-phase with similar amplitude, usually heteroclinic [6, 16]. This definition is consistent with the Red Queen hypothesis inspired by Lewis Carroll’s Through The Looking Glass: “it takes all the running you can do” (dominance switching represented by out-of-phase cycles), “to keep in the same place” (oscillations with similar amplitude, representing host fitness remaining the same even though there is coevolution) [16, 20]. Moreover, one of the properties of the canonical Red Queen dynamics is that the population sizes of a host type can reach its maximum near the value of the carrying capacity, but with associated risk of extinction (impermanent coexistence) when negative-frequency selection occurs in favor of other host types [6, 16, 21, 22].

The Red Queen dynamics is akin to the kill-the-winner hypothesis in bacteria-phage systems [23, 24]. The fitness of a common host type decreases due to parasitism, initiating the escalation in the fitness of a rare host type. The new common host type will then be the target of parasitism [6]. When hosts fail to survive the co-evolution, then their population may converge to extinction. Similarly, parasites that fail to catch up with the evolving hosts may soon be eradicated [9]. To simulate the Red Queen dynamics with perpetual negative frequency-dependent selection without extinction, we assume that the persistence of Red Queen dynamics materializes in conditions where hosts and parasite can avoid extinction and recover when they reach very low density [16]. The parameter range leading to the Red Queen dynamics is wide in deterministic host-parasite systems because there is symmetrical selection, that is, host evolution is countered by the evolution of parasites, especially in the absence of alternative hosts [6, 9]. The factors that foster the Red Queen cycles are strong repression to express multiple host types (inter-host type competition), adequate basal host birth rate for survival, sufficient carrying capacity of the host’s environment, high degree of parasite specificity (e.g., matching-allele interactions), and intermediate levels of parasite mortality [16]. Inter-host type competition characterizes evolutionary selection among host types (genotype or phenotype) that will be commonly expressed in the host population. In multi-type host-parasite systems, multiple host types may coexist, but if canonical Red Queen dynamics arises, only one host type is common/dominant for a certain period of time [25].

Intermediate level of parasite mortality is essential in maintaining Red Queen dynamics. Low parasite death rate imposes high degree of parasitism that adversely affects the host populations; while high parasite death rate limits the antagonistic influence of parasites to initiate negative frequency-dependent selection in hosts [16]. If it is desired to escape the coevolution in Red Queen dynamics, then we can do various strategies, such as suppressing the parasitic functional response in host population (e.g., through introduction of probiotics), suppressing the numerical response in parasite population (e.g., through introduction of antibiotics), and increasing death rate of parasites (e.g., through introduction of a different kind of antibiotics). Here, “escape” means stopping the parasitism-driven Red Queen cycles, where the outcome is a surviving stable host population. We focus on the mathematical investigation of the effect of suppressing the numerical response in parasite population through the introduction of antibiotics to attain a stable positive host population abundance. The inhibiting factor that suppresses the growth of parasites is referred to as *antibiotic*; although, our results can be applied beyond bacterial parasites but without side effect to the host.

Our minimal mathematical model, for the first time, explain a strategy for escaping the Red Queen dynamics via application of antibiotics. We considered the conditions favorable in steering the Red Queen dynamics as the starting point of our simulations. Then we show the levels of antibiotic effectivity that can stop the cycles. We also show if discontinuing the application, or even just intermittent intake, of antibiotics will provide opportunity for the parasites to recover. Without losing generality, our model assumes that hosts are killed or castrated once infected by the parasite, that is, they cannot recover from disease nor further reproduce. This is often the case in ecological systems with castrating parasites [26, 27, 28].

## Mathematical Model

Mathematical modeling is very useful in understanding host-parasite interaction and diseases [29, 30, 6, 31]. The phenotype decision-switch network in Figure 1 is used to illustrate the interaction among host phenotypes and parasites, and the effect of antibiotics [25]. Here, the expression of host phenotype 1 (H1) inhibits the expression of host phenotype 2 (H2), and vice versa. Moreover, parasites decrease the frequency of the phenotype they are attacking due to infection. As indicated in the network (Figure 1), parasitism has high specificity, where parasite 1 (P1) and parasite 2 (P2) target H1 and H2, respectively. We assume that there is differential effect of antibiotic to each parasite, and the antibiotic does not have side-effect to any of the hosts. In our simulations, the mathematical model of host-parasite interaction with antibiotic is as follows.

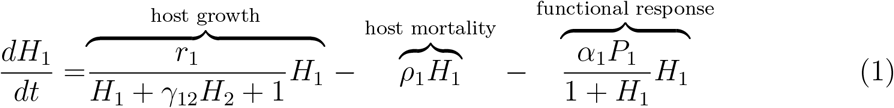

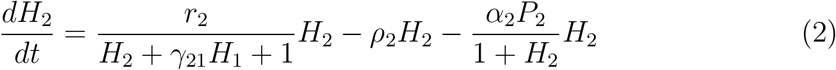

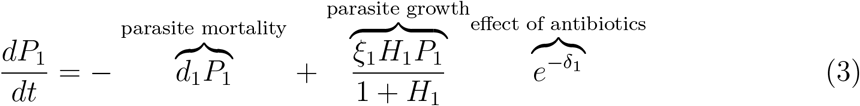

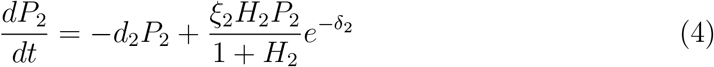

Two host and two parasite types are considered in this study. To highlight the interaction of host and parasite with the effect of antibiotics, certain simplifying assumptions are considered. First, the hosts have similar characteristics, that is, the growth coefficients of H1 and H2 are equal (*r*_1_ = *r*_2_ = *r*), and they have equal death rates (*ρ*_1_ = *ρ*_2_ = *ρ*). Both the parasite growth and death rates are also equal, that is, *ξ*_1_ = *ξ*_2_ = *ξ* and *d*_1_ = *d*_2_ = *d*, respectively. Furthermore, parasitism efficiency is the same for both parasites (*α*_1_ = *α*_2_ = *α*), and effect of antibiotics to both parasites are equal (*δ*_1_ = *δ*_2_ = *δ*). Lastly, the strength of competition for both H1 and H2 is equal to 1 (γ_12_ = γ_21_ = 1), meaning both hosts have similar competitive abilities. These simplifying assumptions characterize a host-parasite system where the hosts are from the same family of species, and the parasites are of closely similar types except that each parasite strain targets different host phenotype. The specificity of parasites characterize a system where the hosts are able to evolve or adapt against the infecting parasite strain. Table 1 summarizes the parameters used in the model and their description.

**Table 1:** Table of parameters. All parameters are non-negative.

| Parameter | Definition |
| --- | --- |
| $r_i$ | Growth coefficient of host $i$ |
| $\rho_i$ | Death rate of host $i$ |
| $\xi_i$ | Numerical response coefficient of parasite growth $i$ |
| $d_i$ | Death rate of parasite $i$ not due to antibiotics |
| $\alpha_i$ | Parasitism efficiency of parasite $i$ in infecting host $i$ |
| $\gamma_{ij}$ | Coefficient associated with the inhibition of host phenotype $i$ by phenotype $j \neq i$ |
| $\delta_i$ | Level of effectiveness of antibiotics targeting the growth rate of parasite $i$ |

All the combinations of parameter values in the simulations (*r*, *ρ*, *ξ* and *d*) conform to the Red Queen dynamics in host-parasite system with Type II functional response [16]. The parameter values presented in Table 2 are used in the simulations (100k simulation runs) shown in Figures 2-5. The tested level of effectiveness of the antibiotics are presented in Table 3. The level of effectiveness is a function of the amount of antibiotic (*Y*) and efficacy (*ϕ*). Without losing generality, we set *δ* = *ϕY*. With the addition of antibiotics, we inspect if the host-parasite coevolution will remain or escape the Red Queen dynamics.

**Table 2:** Table of parameter values used in the simulations. L, ML, M, MH and H respectively mean low, medium low, medium, medium high and high values in relation to the parameter range of the Red Queen dynamics.

| Parameter | L-a | L-b | ML-a | ML-b | M-a | M-b | MH-a | MH-b | H-a | H-b |
| --- | --- | --- | --- | --- | --- | --- | --- | --- | --- | --- |
| $r$ | 1 | 1.5 | 2 | 2.5 | 3 | 3.5 | 4 | 4.5 | 5 | 5.5 |
| $\rho$ | 0.05 | 0.055 | 0.06 | 0.07 | 0.08 | 0.09 | 0.1 | 0.11 | 0.12 | 0.13 |
| $\xi$ | 0.5 | 0.75 | 1 | 1.5 | 2 | 3.5 | 5 | 7.5 | 10 | 12 |
| $d$ | 0.05 | 0.15 | 0.3 | 0.4 | 0.5 | 0.65 | 0.8 | 0.9 | 1 | 1.1 |

**Table 3:** Tested level of effectiveness of the antibiotics

| $\delta$ | $e^{-\delta}$ | $Effectiveness = 1 - e^{-\delta}$ |
| --- | --- | --- |
| 1 | 0.368 | 0.632 |
| 1.5 | 0.223 | 0.777 |
| 2 | 0.135 | 0.865 |
| 2.5 | 0.082 | 0.918 |
| 3 | 0.050 | 0.950 |
| 3.5 | 0.030 | 0.970 |
| 4 | 0.018 | 0.982 |
| 4.5 | 0.011 | 0.989 |
| 5 | 0.007 | 0.993 |
| 5.5 | 0.004 | 0.996 |

Moreover, we simulate cases where antibiotic intake is not continuous. We denote *τ* as the gap period between antibiotic intake. For example, *τ* = 30 means that antibiotics are ingested by the host every after 30 simulation periods. The model that we use to reflect the effect of gaps between antibiotic intakes to the parasite population growth (*i* = 1, 2) is

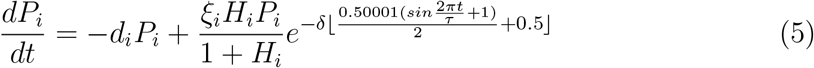

## Results and Discussion

The existence of Red Queen dynamics in host-parasite interaction follows from the existence of parasites, and eradication of parasites can lead to the escape from Red Queen dynamics. We started with an initial state where the host-parasite interaction exhibits Red Queen dynamics. Then we answer the question: what level of antibiotic effectiveness can stop the Red Queen cycles leading to the stable survival of one or more host types? We intend to escape the Red Queen dynamics due to two main reasons: (i) to eliminate or minimize parasite infection leading to the winnerless coevolution of hosts and parasites, and (ii) to avoid the risk of extinction of hosts (impermanent coexistence), especially when there is demographic stochastic noise in host population growth [32]. The possibility of impermanent coexistence in hosts makes the Red Queen dynamics hardly observable in nature [33, 34].

Based on our simulations, the host-parasite interaction still exhibits Red Queen dynamics (Figure 2A) even with the introduction of antibiotics but with low effectiveness. This suggests that the level of antibiotics is inadequate to eradicate parasitism. In the presence of ineffective antibiotics, cyclic host population abundance can still have high amplitude similar to the host-parasite system without antibiotics (Figure 2A). One of the positive consequences of this ineffective antibiotics is that it is possible that the time period before reaching impermanent coexistence widens. However, still the hosts are not able to escape the Red Queen dynamics, with impermanent coexistence as a possible associated recurring risk.

Increasing the level of effectiveness of the antibiotics may lead to a phenomenon wherein the oscillations become irregular (referred to as the edge between the Red Queen dynamics and equilibrium; Figure 2B). This happens when the antibiotics have partial effect on the parasite, driving a decrease in parasite’s efficiency but only for a limited duration. To totally eradicate parasitism and escape the Red Queen dynamics, a greater level of effectiveness of the antibiotics is needed (Figure 2C). The eradication of parasites leads the hosts to a state of equilibrium wherein one or both host types converge to have positive population frequencies. As seen in Figure 2C, both the host types may coexist; however, a specific host could be more dominant than the other host.

As illustrated in Figure 3, the host needs the level of effectiveness of antibiotics to be *δ* > 4 or *δ* > 4.5 in order to escape the Red Queen. Based on Table 3, this indicates that the antibiotic should be > 98.2% or > 98.9% effective to destroy parasites, which is close to an ideal antibiotic. A counter-intuitive result shows that a more effective antibiotic (> 98.9%) is needed to escape the Red Queen dynamics when host growth rate is higher (comparing *δ* associated with *r* < 2.5 and *r* > 2.5 in Figure 3A). This is because a high host growth rate is favorable to generate Red Queen cycles [16].

**Figure 3:**
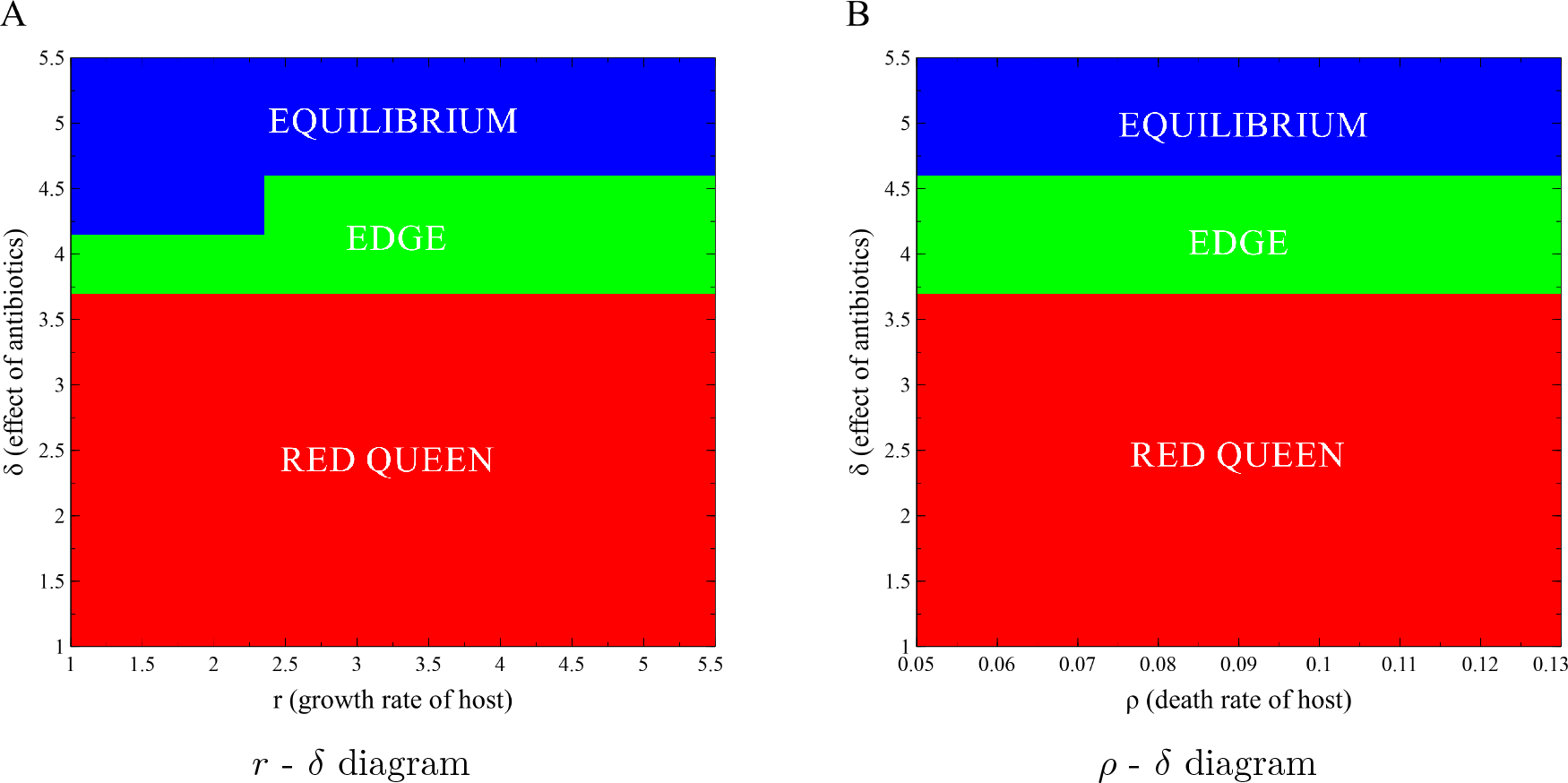
Parameter diagrams showing the qualitative behavior of host populations when varying different phenotypic properties and the level of effectiveness of the antibiotics *δ*. Parameters *d* = 0.05 and *ξ* = 5 are fixed. (A) Varying growth coefficients of hosts (*r*) vs. effectiveness of antibiotics (*δ*); *ρ* = 0.12. (B) Different death coefficients of hosts (*ρ*) vs. effectiveness of the antibiotics (*δ*); *r* = 4.

We investigated the relationship between the parasite growth and death rate, and the effectiveness of the antibiotics. As shown in Figure 4A, the growth rate of the parasite (*ξ*) has a direct relationship to the effectiveness of the antibiotics needed to escape the Red Queen dynamics. The higher the parasite growth rate, the greater the effectiveness of the antibiotics is needed. This is logical since eradication of parasites is more difficult if they have high growth rate. The parasite with less growth rate (*δ* = 1) will be eradicated using a 91.8% effective antibiotic. However, the parasite with high growth rate (*δ* = 10) will be eradicated using 99.6% effective antibiotics. On the other hand, the parasite death rate (*ρ*) has an inverse relationship to the effectiveness of the antibiotics (Figure 4B). As the death rate of the parasite increases, the lower the antibiotic effectiveness is needed to escape the Red Queen dynamics. This is because high parasite mortality (not due to antibiotics) can hasten the parasite’s extermination when antibiotics are introduced. A parasite death rate of *d* = 1 will result in full parasite eradication if we introduce 86.5% effective antibiotics.

Our simulations also show that uninterrupted intake of effective antibiotics could lead to the full eradication of the parasite. However, this result may change when we vary the antibiotic intake of the hosts by taking antibiotics intermittently (i.e., on specific simulation periods only). We investigated the dynamics of the hosts and parasites using the extended model in Equations 5 with varying values of the parameter *τ* with other parameters being fixed. Compared to the case where the antibiotic intake is uninterrupted (Figure 5A), the abundances of both host and parasite populations are low when antibiotics are taken periodically (Figures 5B and 5C). Moreover, the periodic intake of the antibiotics may lead to oscillatory population frequencies with the population dynamics of the hosts having very low amplitude (Figures 5B and 5C). This intermittent intake is more adverse to the hosts compared when the antibiotic is less effective but taken continuously. This suggests that unsustainable antibiotic intake could result in more potent parasite infection. In Figures 5D and 5E, we show that discontinuing the intake of antibiotics, even for a short time, may result in death of hosts. The parasites could become more potent that drives host extinction. This is worse compared to the Red Queen dynamics without antibiotics, since host populations undergoing Red Queen cycles still have the opportunity to survive through deterministic coevolution. Uninterrupted intake of antibiotics is recommended until parasite population is completely eradicated or, possibly, until a level where host immunity can handle the repression of parasite infection.

**Figure 4:**
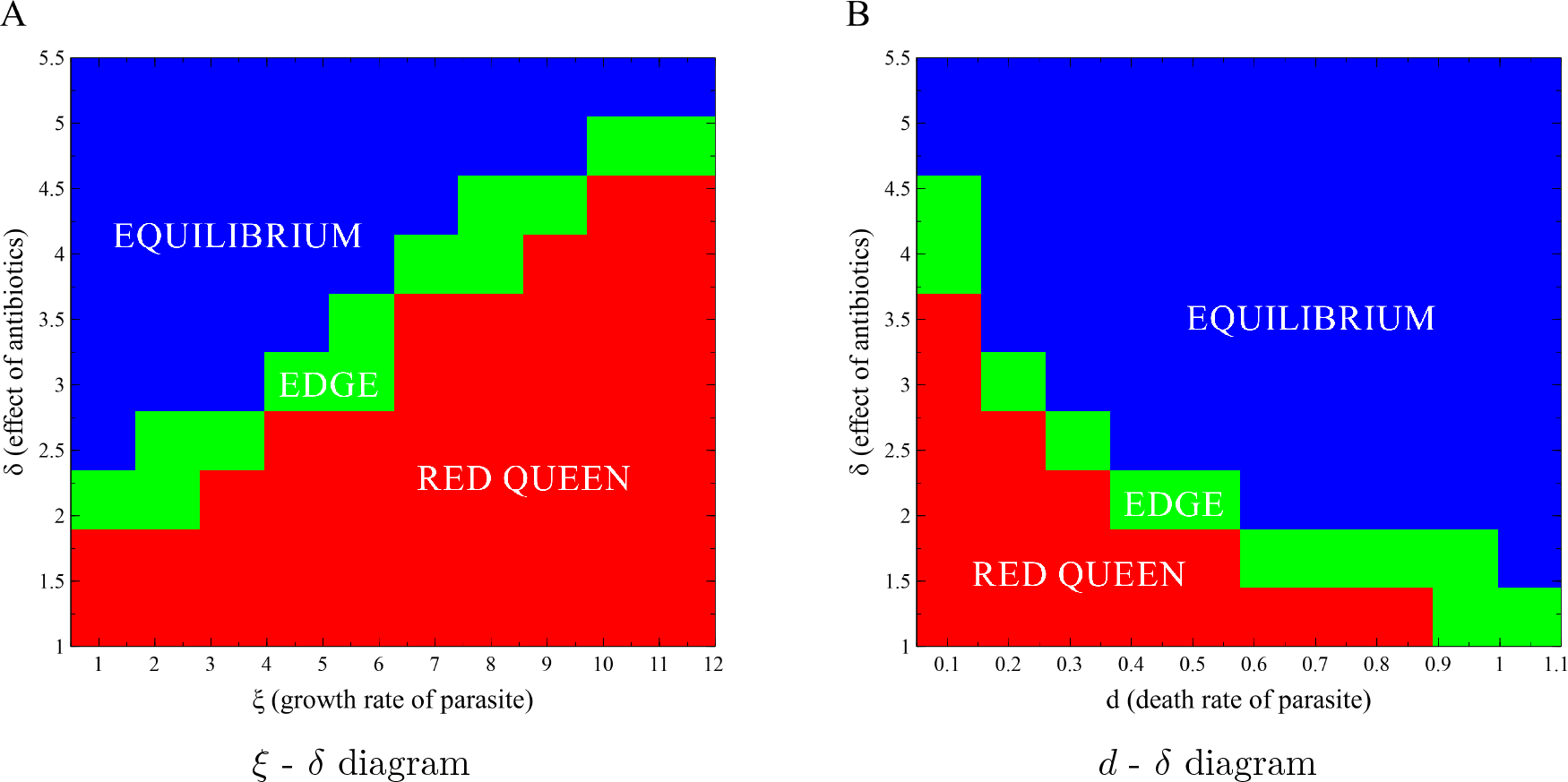
Parameter diagrams showing the qualitative behavior of host populations when varying different parasite characteristics and the level of effectiveness of the antibiotics *δ*. Parameters *r* = 4 and *ρ* = 0.08 are fixed. (A) Parasite growth rates (*ξ*) vs. different levels of effectiveness of antibiotics (*δ*); *d* = 0.05. (B) Death rates of parasites (*d*) vs. different levels of effectiveness of antibiotics (*δ*); *ξ* = 5.

**Figure 5:**
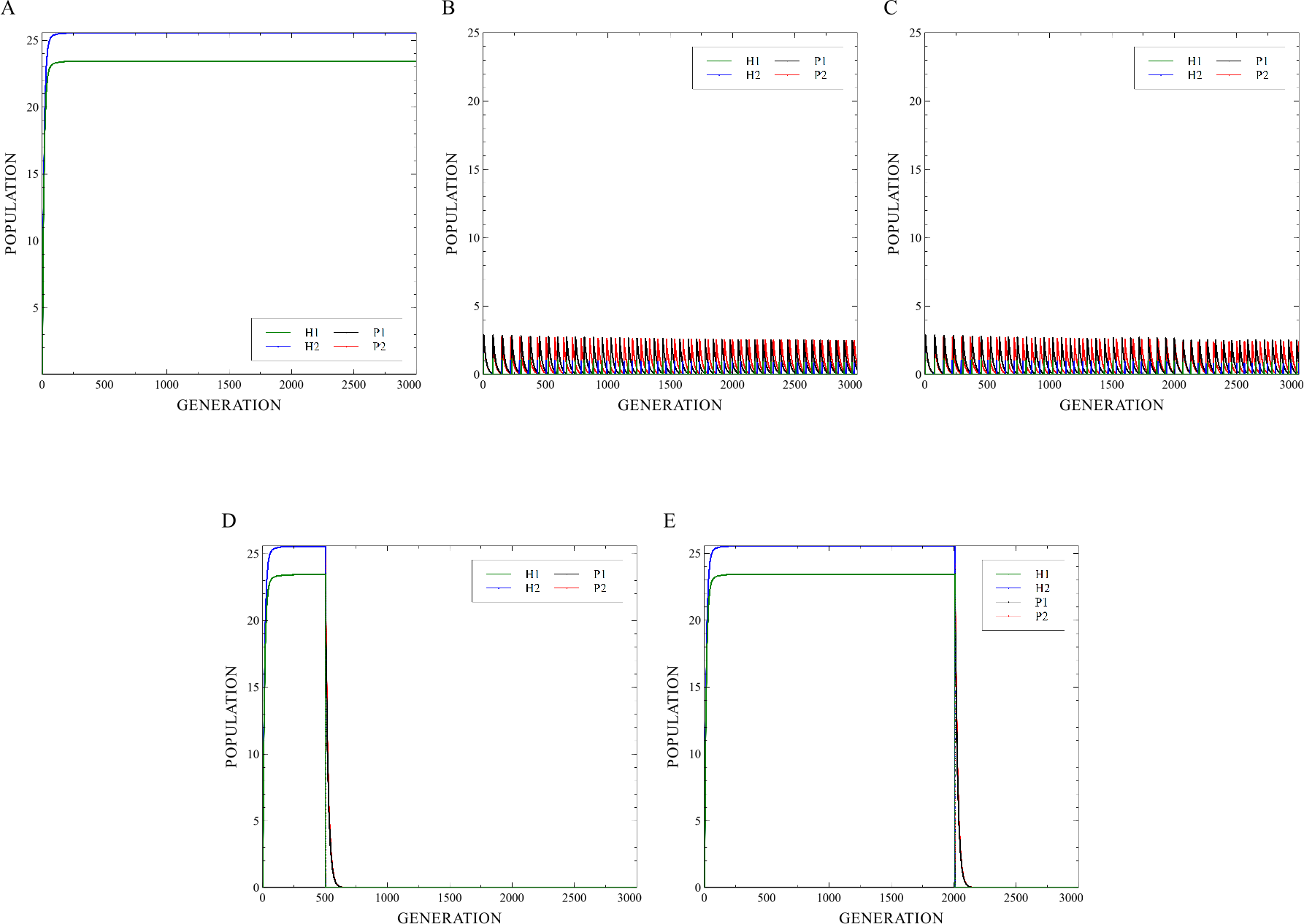
Different behavior of antibiotic intake. (A) Equilibrium convergence showing the coexistence of the hosts and the complete eradication of the parasites because of the uninterrupted intake of antibiotics. (B) Oscillating population of the hosts and parasites but with very low amplitude; the intake of antibiotic is every 1 simulation period. (C) Oscillating population of the hosts and parasites but with very low amplitude; the intake of antibiotic is every 30 simulation period. (D) Equilibrium dynamics with a sudden drop in host population abundance when the antibiotic was taken continuously but is discontinued after 500 simulation periods. (E) Equilibrium dynamics with a sudden drop in host population abundance when the antibiotic was taken continuously but is discontinued after 2000 simulation periods.

## Conclusion

Using mathematical simulations, we show two novel testable hypotheses: (i) for the hosts to escape the Red Queen dynamics, a high level of antibiotic effectiveness is needed; and (ii) intermittent or discontinued intake of antibiotic could be detrimental to the hosts. Here, the introduced antibiotics suppress the growth of the parasites which will then result in parasite eradication. However, if the intake of the antibiotics is discontinued, surviving parasites can recover and these parasites can be more potent causing sudden extinction of hosts. This suggests that for the hosts, the case where antibiotics are introduced but cannot be sustained is riskier compared to the case of Red Queen dynamics without antibiotics. Red Queen dynamics can drive impermanent coexistence in hosts but the hosts have the chance to survive through deterministic cyclic coevolution. Thus, the use of antibiotics should be regulated ensuring its effectiveness even though costly.

## Data Availability

Code available in Dryad [35].

## Author Contributions

JFR conceived the study. JFR and ELA formulated and analyzed the model. ELA implemented the simulations. JFR and ELA wrote the manuscript. All authors reviewed the manuscript and gave final approval for publication. The authors agree to be accountable for all aspects of the work.

## Acknowledgements/Funding

This work was partly supported by UP System Enhanced Creative Work and Research Grant (ECWRG 2016-1-009).

## Research Ethics

Ethical assessment not required to conduct this research.

## Animal Ethics

Ethical assessment not required to conduct this research.

## Permission to carry out fieldwork

No permissions were required since no fieldwork was conducted.

## References

[1] Hatcher, M.J., Dick, J.T., Dunn, A.M., 2012. Diverse effects of parasites in ecosystems: linking interdependent processes. Frontiers in Ecology and the Environment 10, 186–194. (https://doi.org/10.1890/110016)

[2] Wood, C.L., Byers, J.E., Cottingham, K.L., Altman, I., Donahue, M.J., Blakeslee, A.M.H., 2007. Parasites alter community structure. Proceedings of the National Academy of Sciences 104, 9335–9339. (https://doi.org/10.1073/pnas.0700062104)

[3] Hudson, P.J., Dobson, A.P., Lafferty, K.D., 2006. Is a healthy ecosystem one that is rich in parasites? Trends in Ecology & Evolution 21, 381–385. (https://doi.org/10.1016/j.tree.2006.04.007)

[4] Mouritsen, K.N., Poulin, R., 2002. Parasitism, community structure and biodiversity in intertidal ecosystems. Parasitology 124, S101–S117. (https://doi.org/10.1017/S0031182002001476)

[5] Poulin, R., 1999. The functional importance of parasites in animal communities: many roles at many levels? International Journal for Parasitology 29, 903–914. (https://doi.org/10.1016/S0020-7519(99)00045-4)

[6] Rabajante, J.F., Tubay, J.M., Ito, H., Uehara, T., Kakishima, S., Morita, S., Yoshimura, J., Ebert, D., 2016. Host-parasite Red Queen dynamics with phase-locked rare genotypes. Science Advances 2, e1501548. (https://doi.org/10.1126/sciadv.1501548)

[7] Mougi, A., Iwasa, Y., 2011. Unique coevolutionary dynamics in a predatorprey system. Journal of Theoretical Biology 277, 83–89. (https://doi.org/10.1016/j.jtbi.2011.02.015)

[8] Raffel, T.R., Martin, L.B., Rohr, J.R., 2008. Parasites as predators: unifying natural enemy ecology. Trends in Ecology & Evolution 23, 610–618. (https://doi.org/10.1016/j.tree.2008.06.015)

[9] Brockhurst, M.A., Chapman, T., King, K.C., Mank, J.E., Paterson, S., Hurst, G.D.D., 2014. Running with the Red Queen: the role of biotic conflicts in evolution. Proceedings of the Royal Society B: Biological Sciences 281, 20141382–20141382. (https://doi.org/10.1098/rspb.2014.1382)

[10] Morran, L.T., Schmidt, O.G., Gelarden, I.A., Parrish, R.C., Lively, C.M., 2011. Running with the Red Queen: Host-Parasite Coevolution Selects for Biparental Sex. Science 333, 216–218. (https://doi.org/10.1126/science.1206360)

[11] Howard, R.S., Lively, C.M., 2002. The Ratchet and the Red Queen: the maintenance of sex in parasites: Maintenance of sex in parasites. Journal of Evolutionary Biology 15, 648–656. (https://doi.org/10.1046/j.1420-9101.2002.00415.x)

[12] Weitz, J., Wilhelm, S., 2012. Ocean viruses and their effects on microbial communities and biogeochemical cycles. F1000 Biology Reports 4, 17. (https://doi.org/10.3410/B4-17)

[13] Mouritsen, K.N., Poulin, R., 2005. Parasites boost biodiversity and change animal community structure by trait-mediated indirect effects. Oikos 108, 344–350. (https://doi.org/10.1111/j.0030-1299.2005.13507.x)

[14] Lighten, J., Papadopulos, A.S.T., Mohammed, R.S., Ward, B.J., G. Paterson, I., Baillie, L., Bradbury, I.R., Hendry, A.P., Bentzen, P., van Oosterhout, C., 2017. Evolutionary genetics of immunological supertypes reveals two faces of the Red Queen. Nature Communications 8, 1294. (https://doi.org/10.1038/s41467-017-01183-2)

[15] Bonachela, J.A., Wortel, M.T., Stenseth, N.C., 2017. Eco-evolutionary Red Queen dynamics regulate biodiversity in a metabolite-driven microbial system. Scientific Reports 7, 17655. (https://doi.org/10.1038/s41598-017-17774-4)

[16] Rabajante, J.F., Tubay, J.M., Uehara, T., Morita, S., Ebert, D., Yoshimura, J., 2015. Red Queen dynamics in multi-host and multi-parasite interaction system. Scientific Reports 5, 10004. (https://doi.org/10.1038/srep10004)

[17] Jefferies, J.M.C., Clarke, S.C., Webb, J.S., Kraaijeveld, A.R., 2011. Risk of Red Queen dynamics in pneumococcal vaccine strategy. Trends in Microbiology 19, 377–381. (https://doi.org/10.1016/j.tim.2011.06.001)

[18] Ebert, D., 2008. Hostparasite coevolution: Insights from the Daphniaparasite model system. Current Opinion in Microbiology 11, 290–301. (https://doi.org/10.1016/j.mib.2008.05.012)

[19] Decaestecker, E., Gaba, S., Raeymaekers, J.A.M., Stoks, R., Van Kerckhoven, L., Ebert, D., De Meester, L., 2007. Hostparasite Red Queen dynamics archived in pond sediment. Nature 450, 870–873. (https://doi.org/10.1038/nature06291)

[20] Van Valen, L., 1973. A new evolutionary law. Evolutionary Theory 1, 130.

[21] Lutz, A.F., Risau-Gusman, S., Arenzon, J.J., 2013. Intransitivity and coexistence in four species cyclic games. Journal of Theoretical Biology 317, 286–292. (https://doi.org/10.1016/j.jtbi.2012.10.024)

[22] Huisman, J., Weissing, F.J., 2001. Biological conditions for oscillations and chaos generated by multispecies competition. Ecology 82, 2682–2695. (https://doi.org/10.1890/0012-9658(2001)082[2682:BCFOAC]2.0.CO;2)

[23] Jover, L.F., Cortez, M.H., Weitz, J.S., 2013. Mechanisms of multi-strain coexistence in hostphage systems with nested infection networks. Journal of Theoretical Biology 332, 65–77. (https://doi.org/10.1016/j.jtbi.2013.04.011)

[24] Avrani, S., Schwartz, D.A., Lindell, D., 2012. Virus-host swinging party in the oceans: Incorporating biological complexity into paradigms of antagonistic coexistence. Mobile Genetic Elements 2, 88–95. (https://doi.org/10.4161/mge.20031)

[25] Cortez, M.J.V., Rabajante, J.F., Tubay, J.M., Babierra, A.L., 2017. From epigenetic landscape to phenotypic fitness landscape: Evolutionary effect of pathogens on host traits. Infection, Genetics and Evolution 51, 245–254. (https://doi.org/10.1016/j.meegid.2017.04.006)

[26] Ashby, B., Gupta, S., 2014. Parasitic castration promotes coevolutionary cycling but also imposes a cost on sex. Evolution 68, 2234–2244. (https://doi.org/10.1111/evo.12425)

[27] Polak, M., Luong, L.T., Starmer, W.T., 2007. Parasites physically block host copulation: a potent mechanism of parasite-mediated sexual selection. Behavioral Ecology 18, 952–957. (https://doi.org/10.1093/beheco/arm066)

[28] Ebert, D., Joachim Carius, H., Little, T., Decaestecker, E., 2004. The Evolution of Virulence When Parasites Cause Host Castration and Gigantism. The American Naturalist 164, S19–S32. (https://doi.org/10.1086/424606)

[29] Dasbasi, B., Ozturk, I., 2016. Mathematical modelling of bacterial resistance to multiple antibiotics and immune system response. SpringerPlus 5, 408. (https://doi.org/10.1186/s40064-016-2017-8)

[30] Day, T., Read, A.F., 2016. Does High-Dose Antimicrobial Chemotherapy Prevent the Evolution of Resistance? PLOS Computational Biology 12, e1004689. (https://doi.org/10.1371/journal.pcbi.1004689)

[31] Hermsen, R., Deris, J.B., Hwa, T., 2012. On the rapidity of antibiotic resistance evolution facilitated by a concentration gradient. Proceedings of the National Academy of Sciences 109, 10775–10780. (https://doi.org/10.1073/pnas.1117716109)

[32] Allen, E., 2007. Modeling with It stochastic differential equations, Mathematical modelling–theory and applications. Springer, Dordrecht.

[33] Voje, K.L., Holen, O.H., Liow, L.H., Stenseth, N.C., 2015. The role of biotic forces in driving macroevolution: beyond the Red Queen. Proceedings of the Royal Society B: Biological Sciences 282, 2015–0186. (https://doi.org/10.1098/rspb.2015.0186)

[34] Vermeij, G.J., Roopnarine, P.D., 2013. Reining in the Red Queen: the dynamics of adaptation and extinction reexamined. Paleobiology 39, 560–575. (https://doi.org/10.1666/13009)

[35] Anzia, E.L., Rabajante, J.F., 2018. Data from: Antibiotic-driven Escape of Host in a Parasite-induced Red Queen Dynamics. Dryad Digital Repository. (http://dx.doi.org/10.5061/dryad.3c7pd3g) TEMPORARY REVIEW LINK: http://datadryad.org/review?doi=doi:10.5061/dryad.3c7pd3g

